# BioGeoSyn: a graphical software for reproducible biogeographic analyses and cross-clade synthesis of biogeographic events

**DOI:** 10.64898/2026.07.21.739740

**Authors:** Wei Xu, Yun-He Wu

## Abstract

Historical biogeography increasingly looks beyond one clade at a time, pooling range-evolution and speciation events from lineages that share a region to ask whether the same geological and climatic events shaped them together. Despite the momentum behind this cross-clade synthesis, it still has no dedicated software.
We present BioGeoSyn, an open-source graphical application (with an equivalent scripting interface) that makes a complete, reproducible BioGeoBEARS analysis point-and-click: guided data upload and validation, model fitting and comparison, biogeographic stochastic mapping (BSM), and standardized tables, publication figures and a report.
Its one novel capability is cross-clade synthesis. Once every clade’s events carry the same seven process labels and sit on a common time axis, BioGeoSyn integrates event rates through time, overall and region-resolved, across independently analysed clades, each carrying a 95% interval propagated from the stochastic maps, to reveal shared biogeographic pulses.
BioGeoSyn lowers the barrier to reproducible historical biogeography for non-programmers and provides a ready-to-use route to multi-clade biogeographic-event synthesis.

**Data and code for peer review:** the complete source code, the example data and the worked-example analyses are available to editors and reviewers at https://anonymous.4open.science/r/BioGeoSyn-696B/, an anonymised mirror of the development repository. An archived release is deposited in a public repository; its DOI is withheld here for double-blind review.

## 1. Introduction

Reconstructing where lineages lived in the past, and which processes moved and divided their ranges, is central to biogeography and to macroevolutionary studies of diversification. Parametric range-evolution models, the dispersal–extinction–cladogenesis (DEC) model (Ree and Smith 2008), its founder-event extension DEC+J (Matzke 2014), and DIVA-like and BAYAREA-like variants (Ronquist 1997; Landis et al. 2013), are now routinely fitted with BioGeoBEARS (Matzke 2013, 2014), the de facto standard implementation. Biogeographic stochastic mapping (BSM), an application of stochastic character mapping (Huelsenbeck et al. 2003) to these models, additionally samples explicit event histories along the tree, so that dispersal, extinction, vicariance and founder events can be counted and dated (Dupin et al. 2017).

Increasingly, the most compelling biogeographic questions are posed not for a single clade but across many co-distributed lineages, to ask whether shared Earth-history events left a common imprint. A growing body of work extracts range-evolution events from many independently analysed clades, or from very large trees partitioned into clades, and integrates them: to show that in-situ speciation dominates the assembly of global plant diversity (Daru et al. 2026), that diversification bursts and biotic turnovers in Asian mammals track Cenozoic geoclimatic events (Feijó et al. 2022), that Hengduan Mountains uplift drove synchronous diversification across plant clades (Xing and Ree 2017; Ding et al. 2020), that island geography shaped tropical Asian rattan palms (Kuhnhäuser et al. 2025), and that the Himalayan and Hengduan vertebrate faunas share concentrated Miocene diversification pulses (Xu et al. 2021; Lu et al. 2025). Such community-level inference hinges on expressing every clade’s events in the same categories, on a common time axis, with comparable uncertainty, yet this standardization is today assembled by hand, study by study.

Three gaps therefore recur between a fitted BioGeoBEARS model and a defensible, comparable biological narrative.

i. The accessibility gap. BioGeoBEARS is powerful but is driven entirely through R scripting: users hand-assemble the state space, run objects, model set, BSM and plotting. This is a substantial barrier for empiricists without programming experience, and it makes analyses easy to get subtly wrong and hard to reproduce exactly.
ii. The cross-clade gap. The central questions in regional biogeography are comparative: did co-distributed clades disperse or split in synchrony, responding to the same geological or climatic events? Answering this requires the same event categories, on the same time axis, with comparable uncertainty, for every clade. The studies above show the payoff, but no general tool standardizes and automates it.
iii. The interpretation and reporting gap. BSM records events under terse internal codes (y, s, v, j, d, e, a), and a full analysis chains validation, fitting, comparison, BSM, figures and reporting. Translating the codes into named processes and assembling a standardized, reproducible report is done by hand and differently by different users.

Existing tools address these only partially RASP (Yu et al. 2020) pioneered a graphical front end to several biogeographic methods and is widely used, but it centres on single-analysis ancestral reconstruction and visualization; standardized, reconciled cross-clade event synthesis is not its focus. Here we present BioGeoSyn, which (1) makes a complete BioGeoBEARS analysis point-and-click and reproducible; (2) standardizes and visualizes its output as named biogeographic processes, without altering any count; (3), its novel capability, integrates those events across clades through time and space, with uncertainty propagated from the stochastic maps. BioGeoSyn is an accessibility, standardization, visualization and integration layer over BioGeoBEARS (Table 1). Capability (3) is, to our knowledge, new among biogeographic tools and enables a class of analysis: reproducible, community-level comparison of biogeographic events across many independently analysed clades that previously required custom scripting; the graphical interface (1) is aimed at maximizing uptake among empiricists who do not program.

**Table 1.** Capabilities of BioGeoSyn compared with RASP and direct BioGeoBEARS scripting. ✓ = built-in; ∼ = possible with manual effort or partial support; — = not available. Entries reflect typical, documented use and are not exhaustive; all three ultimately rely on BioGeoBEARS for model fitting.

| Capability | BioGeoSyn | RASP | BioGeoBEARS |
| --- | --- | --- | --- |
| No-code graphical interface | ✓ | ✓ | — |
| Guided input validation and data overview | ✓ | ✓ | — |
| Reproducible, config-driven workflow with manifest | ✓ | ~ | ~ |
| Fits full DEC / DIVALIKE / BAYAREALIKE (±J) set | ✓ | ✓ | ✓ |
| Biogeographic stochastic mapping (BSM) | ✓ | ~ | ✓ |
| Standardized named-process event tables and figures | ✓ | — | ~ |
| Reconciliation of event counts to BioGeoBEARS totals | ✓ | — | — |
| Event rates through time with 95% interval | ✓ | — | ~ |
| Region-resolved event budgets and dispersal routes | ✓ | ~ | ~ |
| Source-to-recipient region exchange matrix | ✓ | — | ~ |
| Cross-clade event integration | ✓ | — | — |
| Automatic report and portable result bundle | ✓ | — | — |

## 2. Description and implementation

### 2.1 Overview and scope

BioGeoSyn is, first, a graphical software: a Shiny interface (Chang et al. 2026) guides a user through an entire BioGeoBEARS analysis without writing code. Every graphical action maps onto an auditable R backend, and the same operations are available as ordinary functions, so scripted and point-and-click analyses are byte-for-byte identical. We emphasize what the package is not: it implements no new model, likelihood or estimator. All inference, model fitting and BSM, is performed by BioGeoBEARS itself; BioGeoSyn prepares inputs, standardizes and visualizes outputs, and integrates them across clades. Every derived table is traceable to, and reconciles with, the raw BioGeoBEARS output (§2.3). BioGeoBEARS is an external dependency and is not bundled; the application checks for it at runtime and records its version and citation.

### 2.2 A guided, reproducible workflow

An analysis is fully specified by a single YAML configuration: the tree, the geographic range matrix, region definitions, the maximum range size, the models to run, and options including time-stratified constraints, which the graphical interface writes for the user. A single call to run_workflow() (or one click in the app), then (1) validates inputs with actionable diagnostics; (2) builds the state space and BioGeoBEARS run objects; (3) fits DEC, DEC+J, DIVALIKE, DIVALIKE+J, BAYAREALIKE and BAYAREALIKE+J, caching completed models by input signature so interrupted runs resume; (4) standardizes raw output into tidy tables; (5) optionally runs BSM; (6) generates publication figures; and (7) renders a Quarto report. Raw BioGeoBEARS objects are kept separate from the derived tables, figures and report, the exact configuration and session information are saved, every output is inventoried in a manifest, and the results are packaged into a portable archive (Fig. 1). The workflow is thus reproducible and auditable by construction. Every run also writes an executable R script, reproduce.R, which reads the saved configuration, repoints it at the copies of the inputs stored with the results, and re-runs the analysis end to end; a recipient of the archive can therefore regenerate every table and figure from the command line without opening the interface.

**Figure 1.**
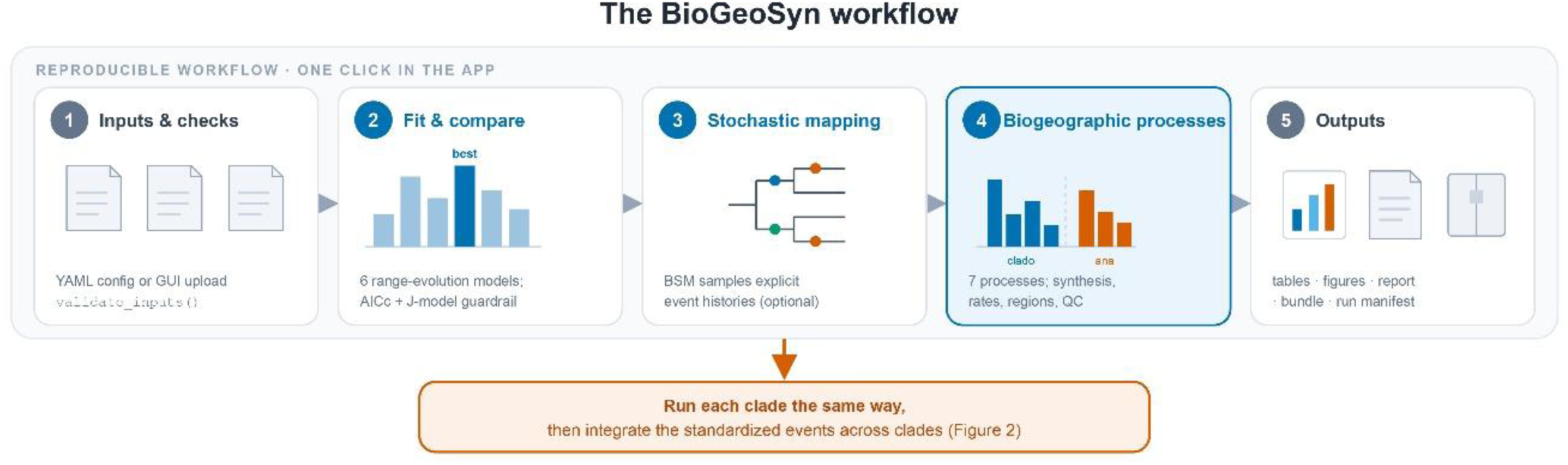
The BioGeoSyn workflow. A single point-and-click run in the Shiny application (equivalently, one run_workflow() call) drives a reproducible five-stage pipeline: (1) input upload, validation and preparation; (2) fitting and comparing six range-evolution models with a guardrail on the founder-event (+J) models; (3) optional biogeographic stochastic mapping (BSM); (4) standardization of BSM output into seven named biogeographic processes, with a per-process synthesis, rates through time (95% intervals), region budgets and dispersal routes, and a BSM reliability check; and (5) portable outputs (standardized tables, publication figures, a Quarto report, a bundle and a run manifest). The top bar denotes the graphical interface; the grey band, the equivalent reproducible workflow. Function names are shown in monospace.

### 2.3 Standardized, interpretable event outputs

To make BSM output interpretable and, critically, comparable across clades, BioGeoSyn relabels BioGeoBEARS’ event codes into a fixed, exhaustive vocabulary of seven named biogeographic processes in two classes (Table 2): cladogenetic speciation modes realized at nodes: in-situ / sympatric speciation (y), subset sympatry (s), vicariance (v) and founder-event / jump speciation (j); and anagenetic range changes along branches: range expansion (d), local extinction (e) and range switching (a). From this, summarize_biogeographic_processes() builds a per-process table of mean counts, standard deviations and proportions, and the package resolves the same events through time (binned mean counts, with a 95% interval taken as the 2.5–97.5% percentiles across stochastic maps) and by region (per-area event budgets and directed dispersal routes), together with a BSM reliability check.

**Table 2.**
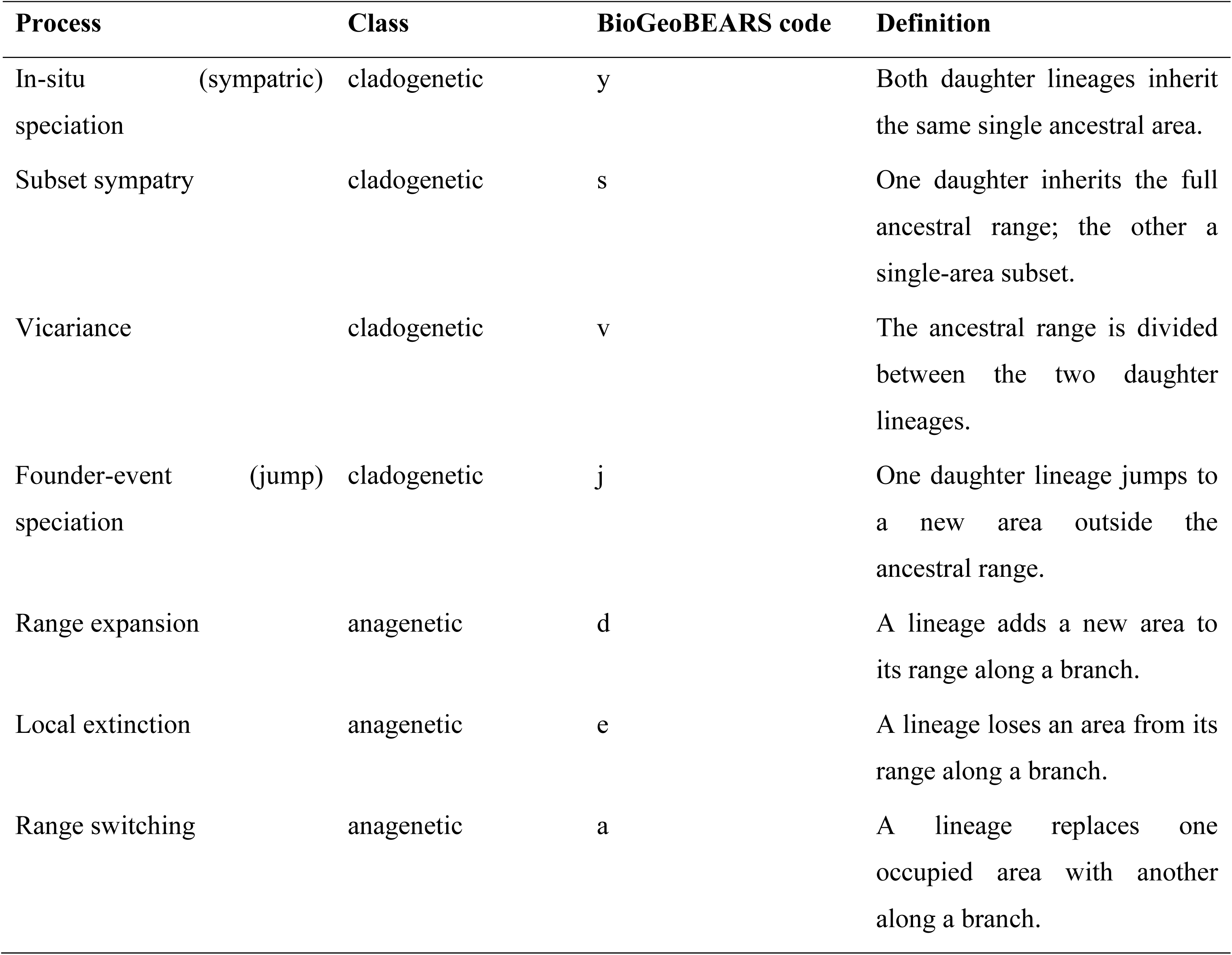
The biogeographic process taxonomy. Mapping from BioGeoBEARS stochastic-mapping event codes to the named biogeographic processes used throughout BioGeoSyn. Generated by biogeographic_process_taxonomy().

This layer is deliberately a standardization, not a new estimator, and we verify that it changes nothing: because the mapping is one-to-one and exhaustive, the four cladogenetic process means sum to BioGeoBEARS’ all-cladogenetic total, the three anagenetic means to the all-anagenetic total, and together to the total event count. This reconciliation is enforced by an automated test, so the standardized tables are guaranteed to represent the underlying counts.

### 2.4 Cross-clade synthesis

Because the events of every clade are expressed in the same named processes on a shared before-present time axis, BioGeoSyn can integrate results across independently analysed clades, the one capability that distinguishes it from existing tools (Fig. 2). A user analyses each clade once with the single-clade workflow (with BSM enabled) and downloads its portable result bundle; the cross-clade tab then takes a batch upload of these bundles and reads the standardized tables from each, so no re-analysis is required and every integrated number remains traceable to a specific clade’s stochastic maps.

**Figure 2.**
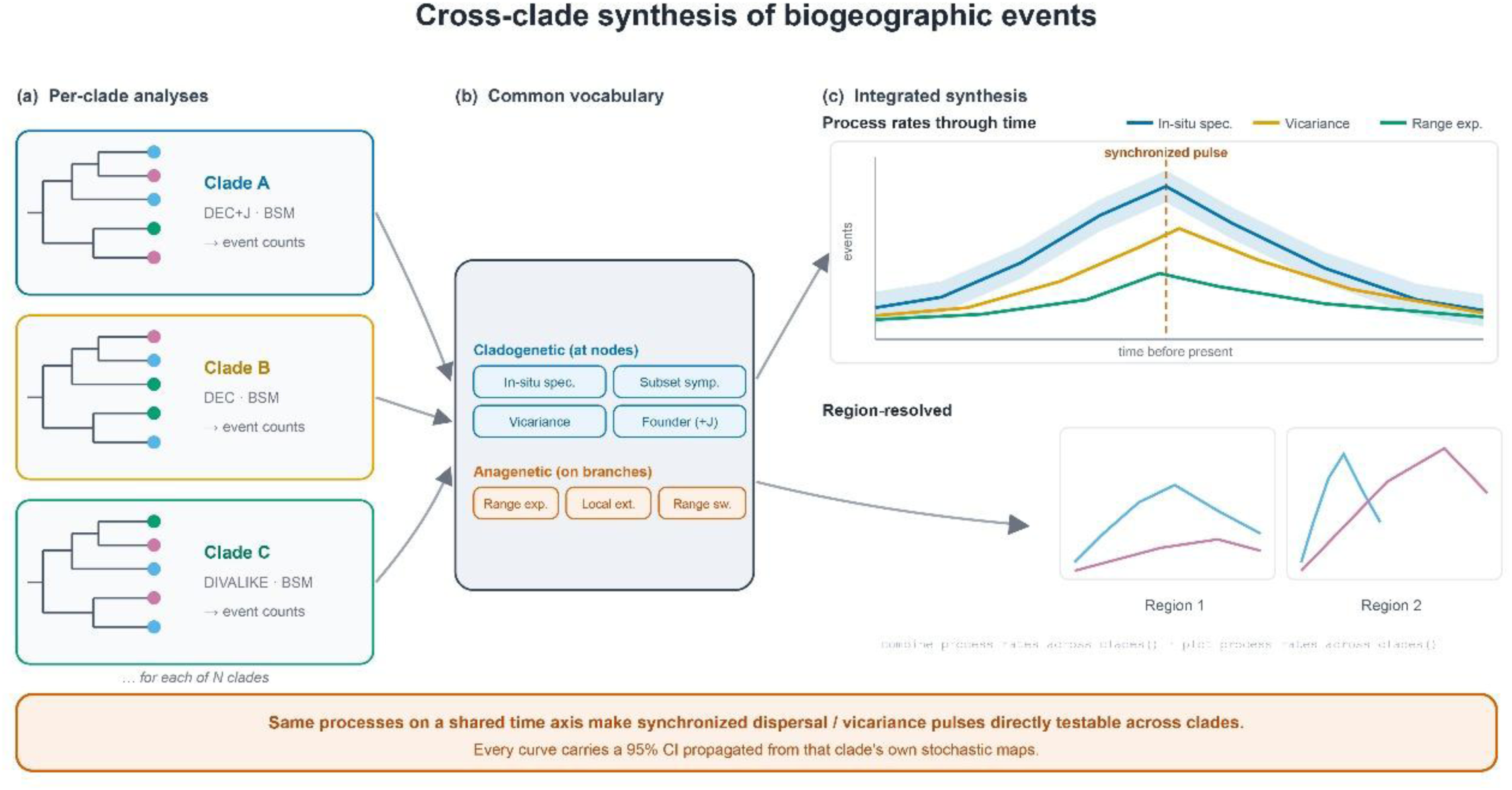
Cross-clade synthesis: the novel capability. Independently analysed clades (a) each yield BioGeoBEARS stochastic-mapping event counts. Because every clade’s events are expressed in the same vocabulary of seven biogeographic processes (four cladogenetic, three anagenetic) on a shared before-present time axis (b), event rates can be integrated across clades (c). Process rates through time (one curve per clade, with a 95% band propagated from each clade’s own stochastic maps) reveal synchronized pulses (dashed line); the region-resolved view pools the clades onto a shared time grid and shows one panel per region with a curve per process, localizing shared signals in space as well as time. This turns per-clade analyses into a community-level, comparative synthesis without altering any clade’s inference.

From the uploaded bundles BioGeoSyn assembles a family of integrated views. combine_process_rates_across_clades() and plot_process_rates_across_clades() draw process rates through time, one panel per process: pooled across clades onto a shared time grid to show the aggregate temporal pulse, or, the same function’s alternative, one curve per clade with a 95% band, so a synchronized burst of, for example, range expansion at a particular time can be examined at either resolution. Pooling matters because independently dated trees never share bin edges; each clade’s native bins are redistributed by overlap onto a common grid so totals are conserved. The region-resolved counterpart applies the same pooling and shows one panel per region with a curve per process (in-situ speciation, immigration, emigration), with the interface exposing the time-bin width, an optional log axis, and a region selector so the user can, for example, restrict the view to the focal regions. BioGeoSyn further integrates a per-process synthesis summed across clades, a source-to-recipient exchange matrix, per-region immigration/emigration budgets, event statistics, and a directed dispersal network between regions that can be sliced by geological period (Paleogene, Neogene, Quaternary, or the total) directly from the pooled event times. Uncertainty is propagated from each clade’s own stochastic maps rather than assumed, and clades need only share comparable time units and region definitions. The tab exports every integrated table and figure as one bundle and can render a self-contained HTML report of the whole synthesis.

This turns per-clade BioGeoBEARS analyses into the building blocks of a community-level, comparative synthesis: a direct, reproducible way to test hypotheses of concerted dispersal or vicariance, while leaving each clade’s inference untouched and fully traceable.

### 2.5 Model-selection guardrails

By default the workflow separates the best-fitting statistical model from biological interpretation. It reports model uncertainty and +J sensitivity and does not auto-declare a single winner; when a +J model is best or near-best, the report and interface flag interpretation cautions rather than asserting founder-event speciation (Ree and Sanmartín 2018; Klaus and Matzke 2020).

### 2.6 Comparison with existing tools

Table 1 places BioGeoSyn alongside RASP and direct BioGeoBEARS scripting across twelve capabilities. All three fit the same model set, so all three can answer the classical single-clade question of where a lineage originated; they differ in what surrounds the fitting step. RASP (Yu et al. 2020) is a mature and widely used graphical environment, and it spans a broader range of inference methods than BioGeoSyn, which targets BioGeoBEARS alone. On the front end the two are comparable: both remove the need to write code, and both guide the user through input preparation and model fitting. The difference lies downstream of the fitted model, where RASP centres on the ancestral reconstruction and its visualization for one analysis at a time. Direct scripting sits at the opposite extreme. It can in principle produce any output discussed here, and experienced users do produce them, but each table and figure has to be written from scratch, and the resulting code is seldom portable between studies. Three capabilities have no counterpart in either alternative. Event counts are reconciled against BioGeoBEARS’ own cladogenetic and anagenetic totals, so every derived table is auditable rather than merely convenient. Each run is packaged as a portable bundle with a manifest and an executable script, so a result can be re-run by someone who did not produce it. And events from independently analysed clades can be integrated in time and space, with uncertainty carried through from each clade’s stochastic maps. The last of these is the capability the rest of the package exists to support.

### 2.7 Use of AI tools

The software and this manuscript were prepared with the assistance of Claude Opus 4.8 (Anthropic), a large language model. The tool was used to help implement and debug the R code of BioGeoSyn, and made no scientific or interpretive contribution. All conceptual and design decisions, including the core idea, the user interface, the software architecture, the data design, and the design of the study and its worked examples, were made by the authors. The authors reviewed and tested all generated code, verified every reported result against the underlying analyses, and reviewed and approved all text. The corresponding author takes full responsibility for all code and text, including any implications for copyright and intellectual property.

## 3. The graphical interface

The application is a five-step wizard aimed squarely at users who do not program (Fig. 3). (1) Data: the user uploads a tree, a geography matrix and a regions table (each with a downloadable template), optionally adds time-stratified constraints, sets the maximum range size and the number of CPU cores, and immediately sees a data overview (tips, species per region, range-size distribution), a state-space size warning, and input validation. (2) Analysis: a single button runs the workflow, with a check-only “dry run” and an optional BSM toggle. (3) Single clade: the reconstruction under the best-fitting model, shown both as probability pies and as a single-most-likely-range view, beside the model comparison and +J caution, with one-click download of the full result bundle. (4) Multi-clade synthesis: the batch-upload integration of §2.4, where the per-clade process synthesis, event statistics, exchange matrix and dispersal network are produced for the pooled clades, together with a downloadable report and bundle. (5) About and citation: version, environment, developer and citation information. Because every step calls the same backend functions used at the command line, the graphical and scripted analyses yield identical results; the interface adds accessibility, not a second code path.

**Figure 3.**
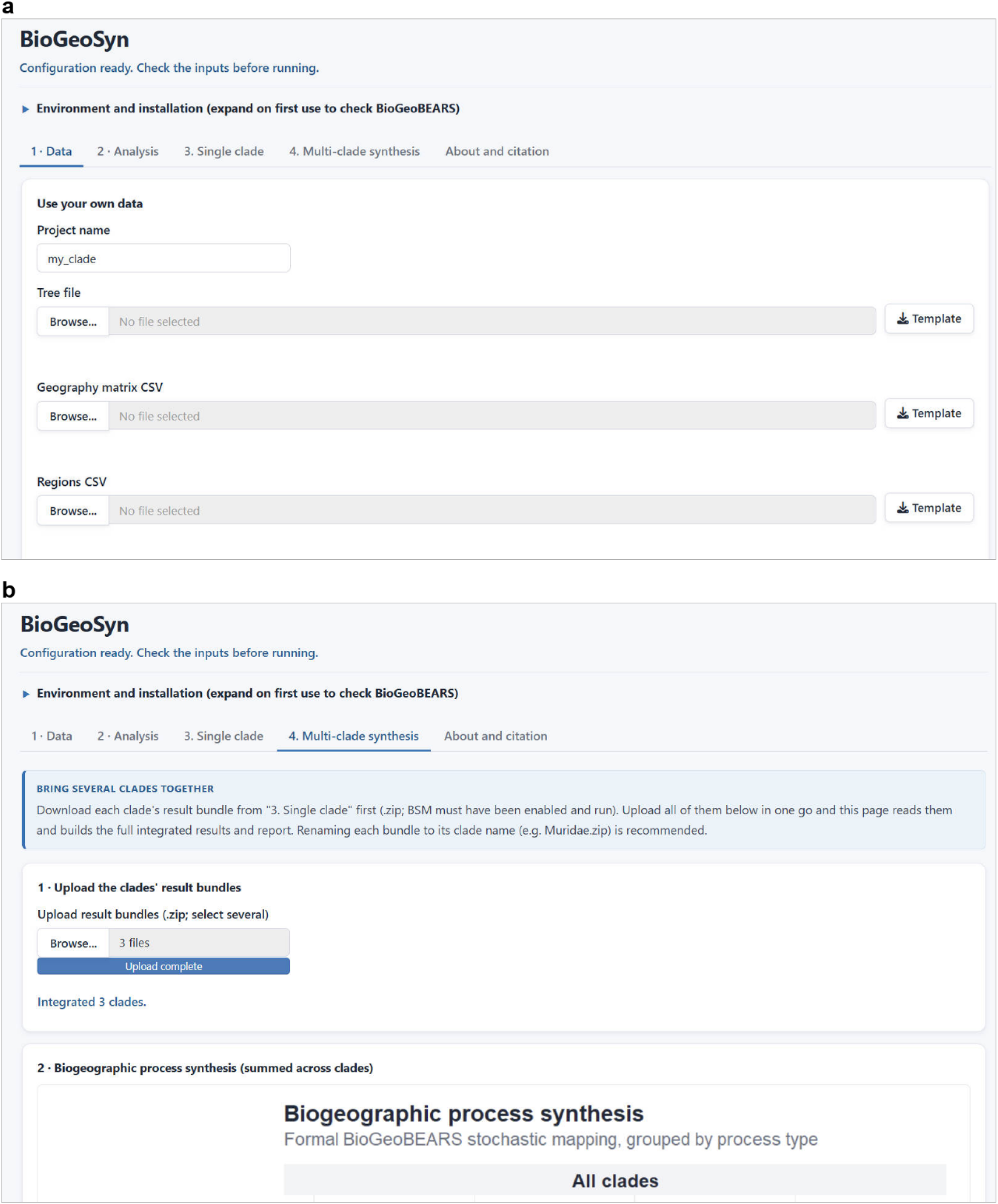
The graphical interface. (a) The data step, where the user supplies a tree, a geography matrix and a regions table, each with a downloadable template, and sees input validation and a data overview before any model is fitted. (b) The multi-clade synthesis step, where the result bundles downloaded from individual single-clade runs are uploaded together in one operation; the interface reports how many clades were read and then assembles the pooled process synthesis, rates through time, region budgets, exchange matrix and period-sliced dispersal networks, together with a downloadable report and bundle.

## 4. Worked example

We illustrate BioGeoSyn at both scales it is designed for: a single-clade reconstruction reproduced against its source study, and the cross-clade synthesis applied to a published multi-clade dataset. In both cases the test is whether the published inferences are recovered.

### 4.1 A single clade: torrent frogs (*Amolops*)

We re-analysed the 23-species phylogeny of the torrent frog genus *Amolops* (Wu et al. 2020) across the four regions those authors defined (the Himalayas, the Hengduan Mountains, Southeast China, and the Indo-Burma/Thai-Malay Peninsula) entirely through the graphical interface. Fitting the six models, BioGeoSyn identified DEC+J as best (Akaike weight 0.50), with DIVALIKE+J effectively tied (0.41) and the non-+J models markedly worse (ΔAICc ≥ 9.7); the interface flagged +J as statistically supported but carrying an interpretation caution rather than asserting founder-event speciation. The reconstruction placed the most recent common ancestor of *Amolops* in the Indo-Burma/Thai-Malay Peninsula (marginal probability 0.38 for that area). These results reproduce the published analysis, which likewise selected DEC+J as best with DIVALIKE+J at an Akaike weight of 0.41, and likewise inferred an Indo-Burma/Thai-Malay origin followed by outward dispersal (Wu et al. 2020), obtained here point-and-click. The match is exactly what the re-present, never re-estimate principle predicts: the numbers are BioGeoBEARS’ own, surfaced through a reproducible interface (Fig. 4).

**Figure 4.**
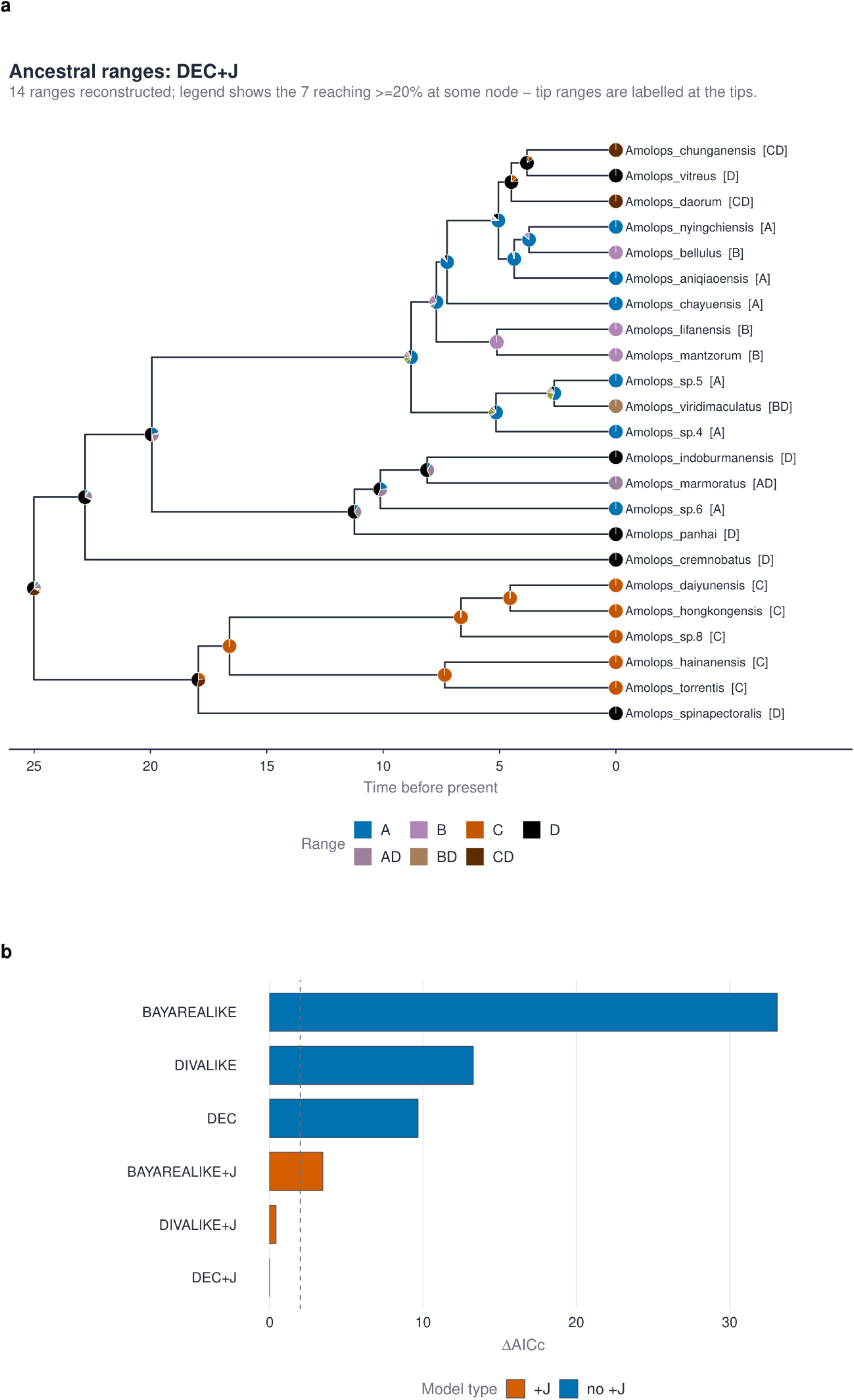
Single-clade worked example: the torrent frogs (Amolops), analysed entirely through the graphical interface. (a) Ancestral ranges under the best-fitting model (DEC+J) for the 23-species phylogeny of Wu et al. (2020), across the four regions those authors defined (A, Himalayas; B, Hengduan Mountains; C, Southeast China; D, Indo-Burma/Thai-Malay Peninsula). Pie charts at internal nodes give the marginal probability of each ancestral range and tip ranges are given in brackets; the most recent common ancestor is reconstructed in the Indo-Burma/Thai-Malay Peninsula. (b) Comparison of the six fitted models as ΔAICc, with founder-event (+J) models distinguished by colour and the dashed line marking ΔAICc = 2. DEC+J is best-fitting, DIVALIKE+J is effectively tied, and every model without +J is more than nine AICc units worse. Both panels are unmodified BioGeoSyn output.

### 4.2 Across clades: Asian mammals

To exercise the novel capability we reused the 21 mammal clades of Feijó et al. (2022) (3,114 species across eleven regions), each analysed with BSM, and uploaded their result bundles to the cross-clade tab. BioGeoSyn pooled the clades onto a shared time axis (Fig. 5a) and recovered the study’s central conclusions. Southern Asia emerges as the dominant source of Asian mammal lineages: it carries by far the largest emigration budget, initiates 28.9% of all between-region dispersals (Feijó et al. reported 27%, “at least three times higher than any other Asian region”), and is the largest net source of lineages, whereas the Himalayas and Hengduan are net sinks, assembled by immigration rather than by in-situ speciation, consistent with their published role as accumulation centres. Slicing the dispersal network by geological period shows dispersal to be minimal in the Paleogene and to burst through the Neogene (Fig. 5b,c), and in-situ speciation is the single largest process overall, again matching the source study. All of these integrated figures and tables (the region-resolved rates, the exchange matrix, the period-sliced networks and the summary report) were produced from the uploaded bundles alone, with uncertainty propagated from each clade’s own stochastic maps. This is precisely the multi-clade integration that such studies have had to assemble with custom scripts, here reduced to a reproducible, point-and-click operation.

**Figure 5.**
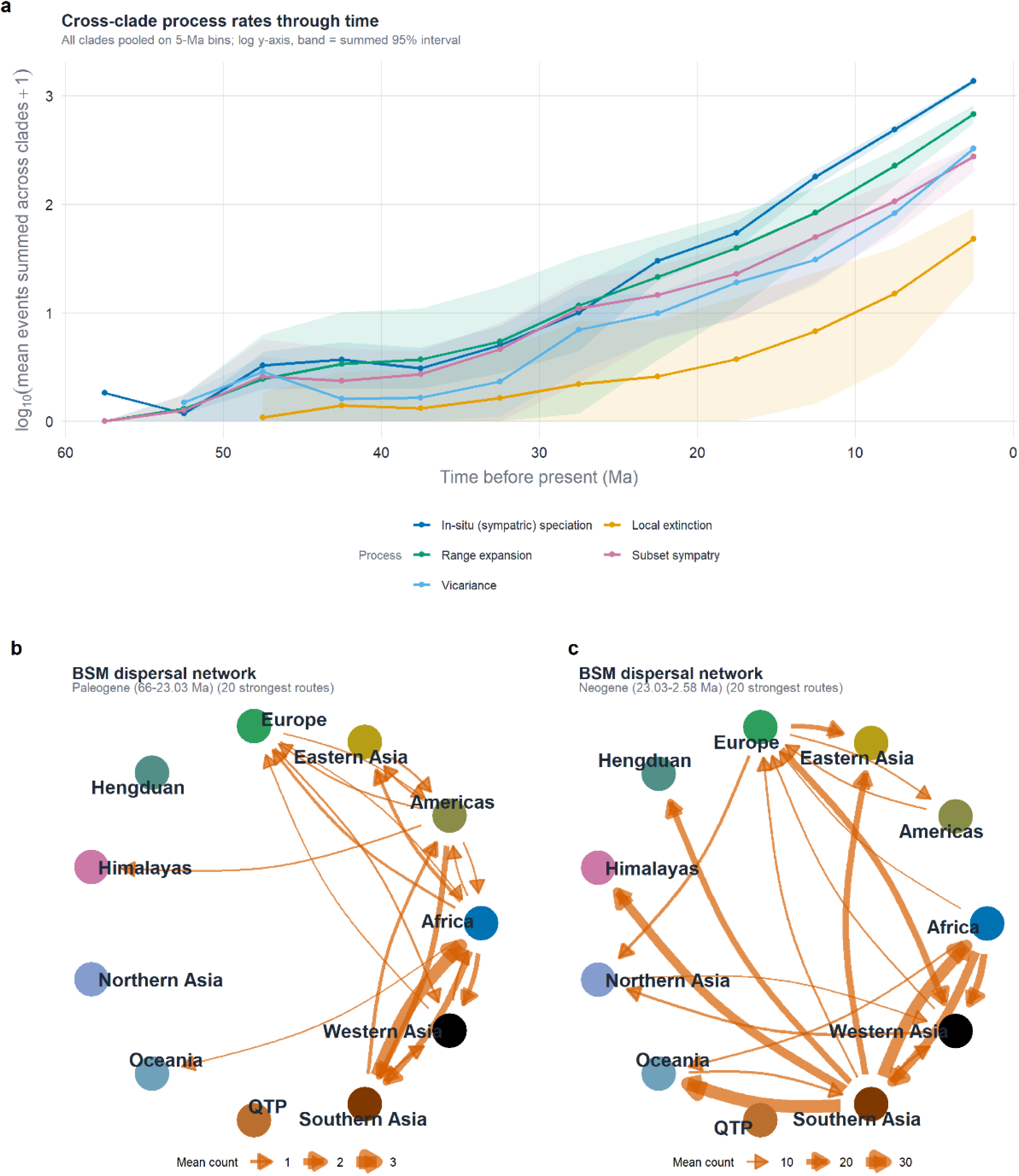
Cross-clade worked example: the 21 Asian mammal clades of Feijóet al. (2022), integrated from their per-clade result bundles with no inference re-run. (a) Process rates through time, pooled across clades onto a shared 5-Ma grid and plotted as log10(mean events + 1); bands are the summed 95% intervals propagated from each clade’s own stochastic maps. Five of the seven standardized processes appear here because every clade was fitted under DEC, which includes neither founder-event (jump) speciation nor range switching. (b, c) Directed dispersal networks between regions for the Paleogene and the Neogene, each showing the 20 strongest routes, with arrow width proportional to the mean number of dispersals per stochastic map. The two scale bars differ by an order of magnitude: dispersal is sparse in the Paleogene and intensifies through the Neogene, with Southern Asia emerging as the dominant source region.

## 5. Discussion

Historical biogeography has, for most of its history, been practiced one clade at a time, and that framing limits what can be concluded. A dispersal pulse or a burst of vicariance recovered in a single group is compatible with many explanations, from a lineage-specific ecological shift to a region-wide reorganization of land and climate, and the observation alone cannot separate them. Only when the same processes are examined across many co-distributed lineages does a shared external driver become testable, because only a common driver is expected to leave a concordant signal in clades that differ in ecology, dispersal ability and age.

The empirical foundations for that comparison are now in place. Time-calibrated phylogenies and curated distributional data exist for a growing number of taxa, and studies that exploit them across clades have become some of the most visible in the field, linking diversification and range change to orogeny, monsoon intensification and Cenozoic climate. What has not kept pace is the analytical infrastructure. Each of those studies had to assemble its own pipeline: event categories were defined locally, time axes were aligned by hand, and the uncertainty carried by stochastic mapping was often discarded on the way to a summary figure. The cost is not only duplicated effort but limited comparability, because two studies that describe the same process under different labels and different binning cannot easily be read against each other. BioGeoSyn is aimed at that bottleneck. By fixing a small, exhaustive vocabulary of biogeographic processes, placing every clade on a shared absolute-time axis, and carrying each clade’s own interval through to the integrated result, it turns cross-clade synthesis from a one-off undertaking into a repeatable operation. Because the standardization is a relabeling rather than a re-estimation, every integrated value remains attributable to the underlying BioGeoBEARS analyses and can be audited against them.

Two limitations follow from this design. BioGeoSyn inherits the assumptions and the sampling of the analyses it summarises, so clades that differ in taxon sampling or in the age of their crown groups contribute unevenly to a pooled curve, and region definitions must be reconciled before clades can be combined at all. It also describes concordance rather than testing it; quantifying whether pulses in different clades are more synchronous than chance expectation is the natural next step, and one the standardized event tables now make straightforward.

## 6. Conclusions

BioGeoSyn makes a complete, reproducible BioGeoBEARS analysis accessible through a graphical interface, standardizes its output as named biogeographic processes, and integrates those events across independently analysed clades in time and space. It adds no new inference: by re-presenting BioGeoBEARS’ own stochastic-mapping output as a named, reconciled vocabulary, it buys interpretability and cross-clade comparability without touching the numbers. Because the graphical interface and the scripting API share a single backend, the same analysis is available to empiricists who do not program and to those who want a fully scripted pipeline. Planned extensions include coupling the cross-clade time axis directly to paleogeographic time-strata.

Availability and requirements

- Source code: https://anonymous.4open.science/r/BioGeoSyn-696B/README.md. Archived release: DOI withheld for double-blind review.
- Programming language: R (≥ 4.1) (R Core Team 2023); figures are built with ggplot2 (Wickham 2016) and phylogenies are handled with ape (Paradis and Schliep 2019). CRAN-hosted dependencies are resolved automatically by the R package manager.
- External dependencies not bundled and not handled by the package manager: BioGeoBEARS, required for analysis, installed separately from GitHub and cited directly (Matzke 2013, 2014); optionally the Quarto command-line tool and a LaTeX engine (e.g. TinyTeX) to render HTML/PDF reports; if either is absent the report step falls back to its source and never blocks a run; and, on Linux, the GLPK and libxml2 system libraries required by igraph.
- Licence: GPL (≥ 2).
- Testing: the package ships an automated testthat suite and is continuously integrated (R CMD check) on Linux, Windows and macOS; a completed code checklist accompanies the submission.
- Reproducibility: every analysis writes an executable R script (reproduce.R) beside the configuration and a copy of the inputs it used, so any result bundle can be re-run from the command line without the graphical interface.
- Coding style: the source follows the tidyverse R style guide (https://style.tidyverse.org/).

## Data availability

The example data distributed with the package, and the worked-example analysis scripts, are archived in the same Zenodo record (DOI: withheld for double-blind review). No new empirical data are generated by this software note.

## Conflict of interest

The authors declare no conflict of interest.

## Statement on inclusion

This study is a software and methods contribution. It involved no new fieldwork, sample collection, or engagement with local communities, and generated no new primary data. The worked examples reuse previously published datasets, and the studies that generated those data are cited in full. The author team includes a researcher based in the region to which the example data relate. As open-source software released under a free licence, BioGeoSyn is intended to be equally usable by researchers in the regions whose biotas are most often the subject of biogeographic study.

## Notes

https://github.com/XuWeiEvo/BioGeoSyn

### Competing Interest Statement

The authors have declared no competing interest.

### Funding

- National Natural Science Foundation of China: NSFC 32300383

